# TDCPP exposure affects the concentrations of thyroid hormones in zebrafish

**DOI:** 10.1101/146092

**Authors:** Jingxin Song

## Abstract

Previous studies show that TDCPP may interrupt the thyroid endocrine system, however, the potential mechanisms involved in these processes were largely unknown. In this study, zebrafish embryos/larvae were exposed to TDCPP until 120 hpf, by which time most of the organs of the larvae have completed development. In this study, the effects of TDCPP on HPT axis were examined and the thyroid hormone levels were measured after TDCPP treatment. Zebrafish (Danio rerio) embryos were treated with a series concentration of TDCPP (10, 20, 40, 80, 160 and 320 μg/L) from 1 day post-fertilization (dpf) to 5 dpf. Exposure concentrations of TDCPP were determined based on the survival rates in each group. Total mRNA were isolated, first-strand cDNA were synthesis and qPCR were performed to detect the mRNA expression levels in hypothalamic-pituitary-thyroid (HPT) axis. The mRNA expression levels of genes involved in thyroid hormone homeostasis were increased in the TDCPP-treated larvae. The mRNA levels of genes involved in thyroid hormone synthesis were also increased in the embryos treated with TDCPP. Furthermore, exposure to TDCPP led to a dose-dependent effect on zebrafish development, including diminished hatching and survival rates, increased malformation. TDCPP treatment significantly reduced the T4 concentration in the 5 dpf zebrafish larvae, but increased the concentration of T3, suggesting the function of thyroid endocrine were interrupted in the TDCPP-exposed zebrafish. Taken together, these data indicated that TDCPP affected the thyroid hormone levels in the zebrafish larvae and could increased the mRNA expression levels of genes related to HPT axis, which further impaired the endocrine homeostasis and thyroid system.

## Background

Tris(2,3-dibromopropyl) phosphate (TDCPP) is an organophosphate flame retardant, which has become one of the most common flame retardant detected in furniture foam [1–5]. During the products’ life cycle, TDCPP releases into the environment due to its unbound feather to the product [5, 6]. TDCPP has been frequently detected in the aquatic system and environment [3, 7–9], including surface water, drinking water, sediments and biota. The toxic effects of TDCPP are largely unclear. Previous studies report that TDCPP adversely affect the neuro-development and interrupt endocrine system [6, 10, 11]. TDCPP can increase the mRNA expression levels of thyroid hormone receptors [6, 11–14].

The hypothalamic-pituitary-thyroid (HPT) axis controls the development of the thyroid endocrine system in fish [15–18], and further regulates the synthesis, secretion and transport of the thyroid hormones[19–22]. Zebrafish is an excellent animal model for environmental monitoring [23–28]. In this study, we aimed to examine the effect of TDCPP on thyroid hormone levels and the mRNA expression levels of genes that modulate HPT axis using zebrafish.

## Materials and Methods

### Chemicals

Tris(1,3-dichloro-2-propyl) phosphate (TDCPP) and 3,3’,5-triiodo-L-thyronine (T3) were purchased from Sigma-aldrich, and dissolved in DMSO as stock solutions.

### Animal maintenance

Adult zebrafish AB line was raised as previous described [29].

### Chemical treatment

Zebrafish were naturally mated and normal eggs were collected for chemical exposure. Fifty 12 hours post-fertilization stage eggs were randomly distributed into 10 cm dishes containing 20 ml different concentrations of TDCPP solutions (0, 10, 20, 40, 80, 160 and 320 μg/L) or 20 μg/L T3 as positive controls. Eggs were treated with drugs until 5 days post-fertilization, and water were changed daily.

### Quantitative real-time PCR

Zebrafish total RNA extraction, cDNA synthesis and qPCR were performed as previously described [30]. Briefly, 50 zebrafish larvae were pooled and homogenized and total RNA were isolated using RNA Isolation Kit (Qiagen). First-strand cDNA was synthesized using RevertAid First Strand cDNA Synthesis Kit (Thermo Fisher Scientific). The qPCR was performed using CFX96 Touch Real-Time PCR Detection System (Bio-Rad). The primers of target genes were described somewhere else [31]. The qPCR program was: 95 °C, 15 min; 40 cycles of 95 °C, 15 s, 60 °C, 1 min. The expression levels of genes were normalized to β-actin using the 2^−ΔΔCt^ method.

### Thyroid hormone measurement

The measurement of thyroid hormone was carried out as described previously [31]. Briefly, 100 zebrafish embryos were homogenized in 0.1 ml ELISA lysis buffer. The samples were centrifuged and supernatants were collected for 3,3’,5-triiodo-L-thyronine (T3) and thyroxin (T4) measurement according to the manufacturer’s instructions (Thermo Fisher Scientific).

### Statistical analysis

The differences between the control and chemical-treated group were examined by one-way analysis of variance (ANOVA) followed by Turkey’s test. P< 0.05 was considered significance.

## Results

### Toxicity of TDCPP to the zebrafish development

Treatment with TDCPP resulted in developmental toxicity to the zebrafish embryos, including lower hatching and survival rates and higher malformation rates (Figures 1–3). In the control group, the survival rate was 85%, and there were no significant differences on the hatching and survival rates in the lower concentrations of TDCPP treated groups (10, 20, 40, 80 and 160 μg/L) at 5 dpf (Figures 1–2). However, the hatching and survival rates were decreased in the 320 μg/L TDCPP treated group compared to the control group at 5 dpf (Figures 1–2). Exposed to 160 and 320 μg/L TDCPP dramatically increased the malformation rates of zebrafish larvae in the 5 dpf, however, exposed to the lower concentrations of TDCPP (10, 20, 40 and 80 μg/L) had no obviously effects on the malformation rates of zebrafish (Figure 3).

**Figure 1.**
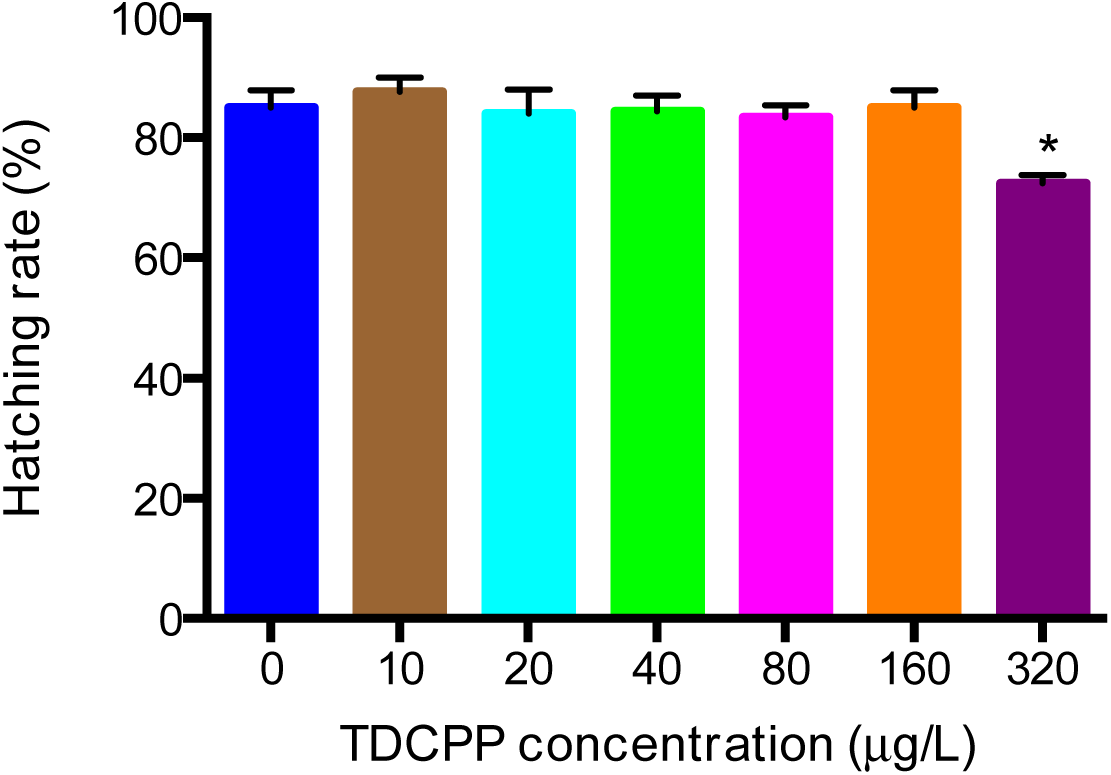
Effects of TDCPP on the hatching rates.

**Figure 2.**
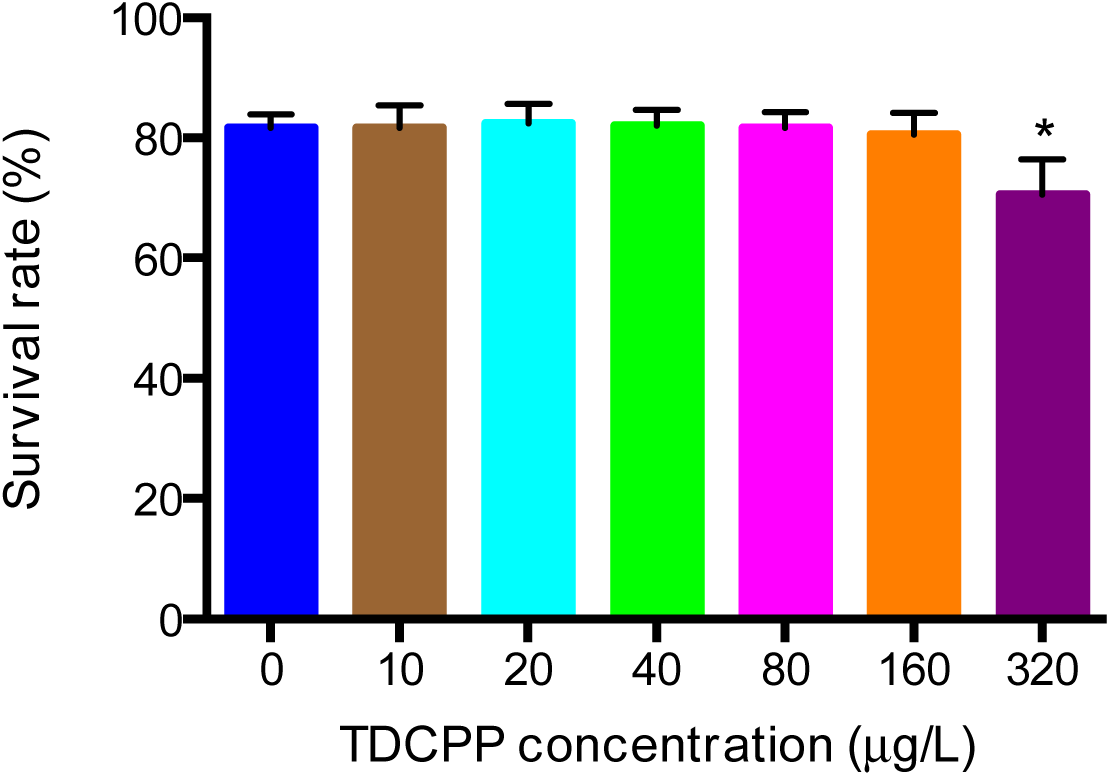
Effects of TDCPP on the survival rates.

**Figure 3.**
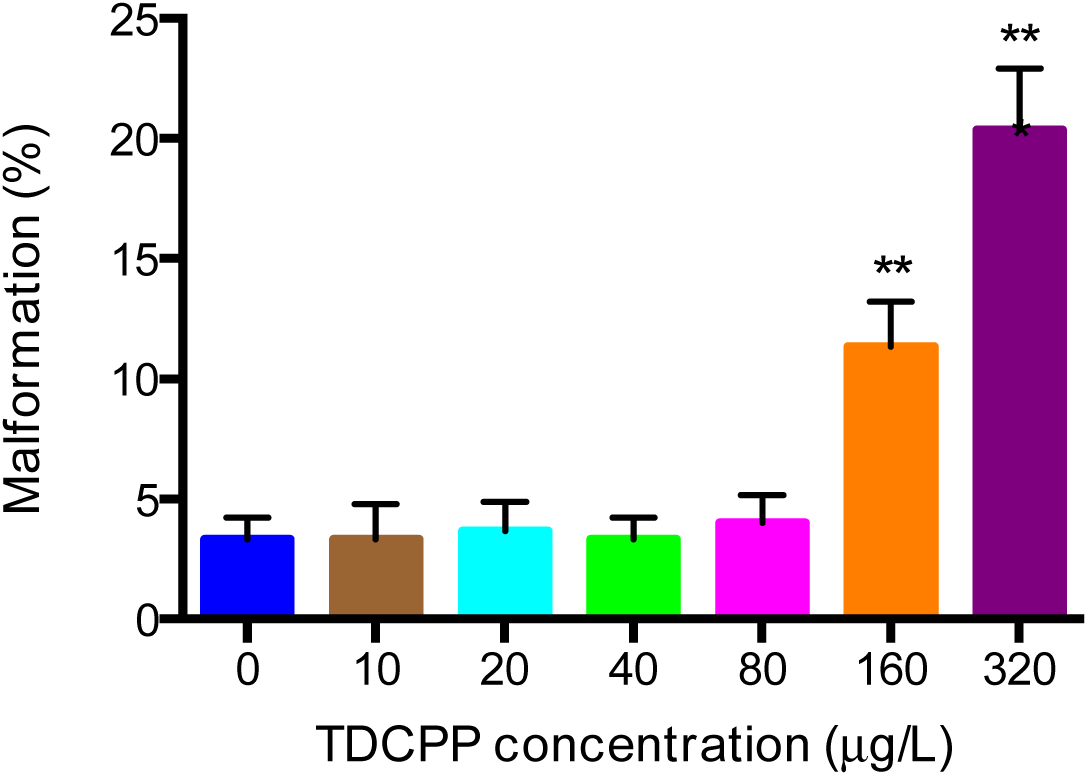
Effects of TDCPP on the malformation.

### TDCPP increased the expression of genes involved in HPT axis

Exposed to T3 or TDCPP affect the mRNA expression levels of genes that were involved in HPT axis in zebrafish development at 5 dpf (Figures 4–5). Exposed to 20 μg/L T3 increased the mRNA expression levels of pax5 and slc5a5, but had no significant effects on nkx2.1, ugt1ab and ttr. There were no obvious changes in the gene levels in the lower concentration treated groups. Exposed to 320 μg/L TDCPP significantly increased all the selected genes expression levels compared to the controls.

**Figure 4.**
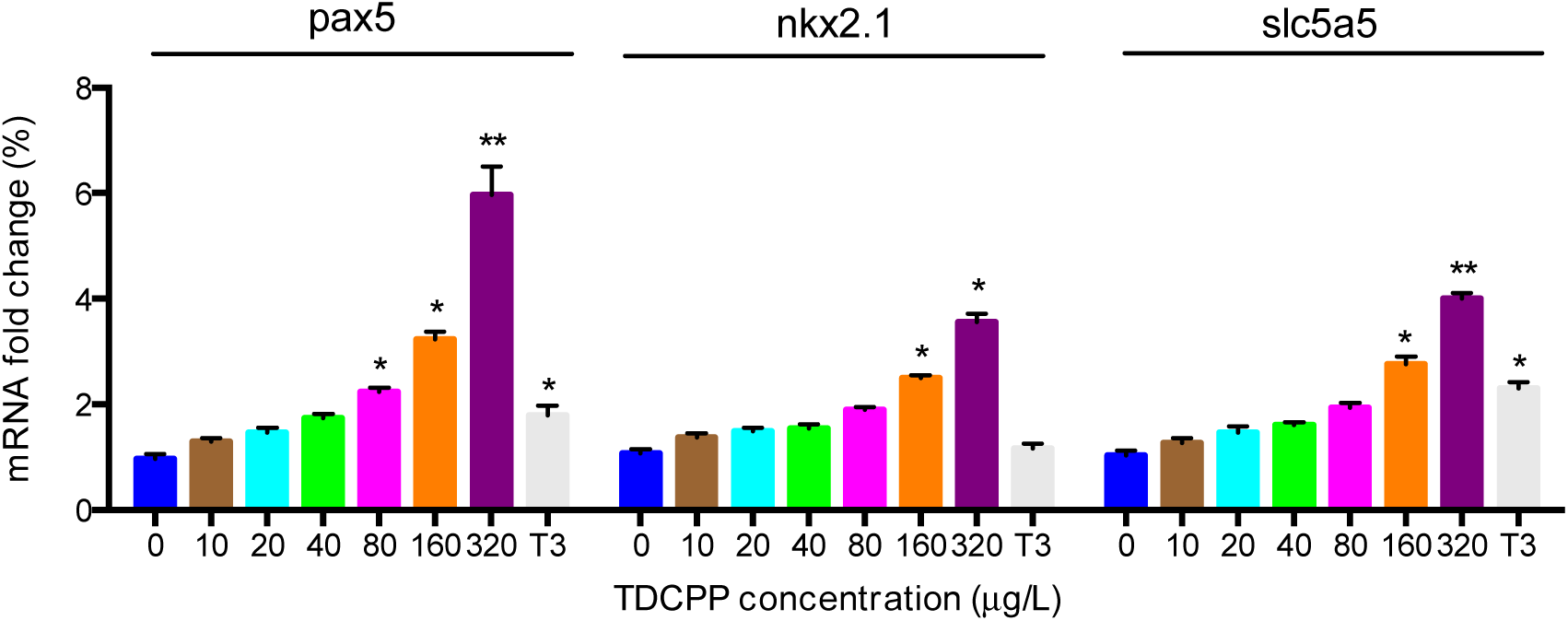
Effects of TDCPP on the expression of pax5, nkx2.1 and slc5a5.

**Figure 5.**
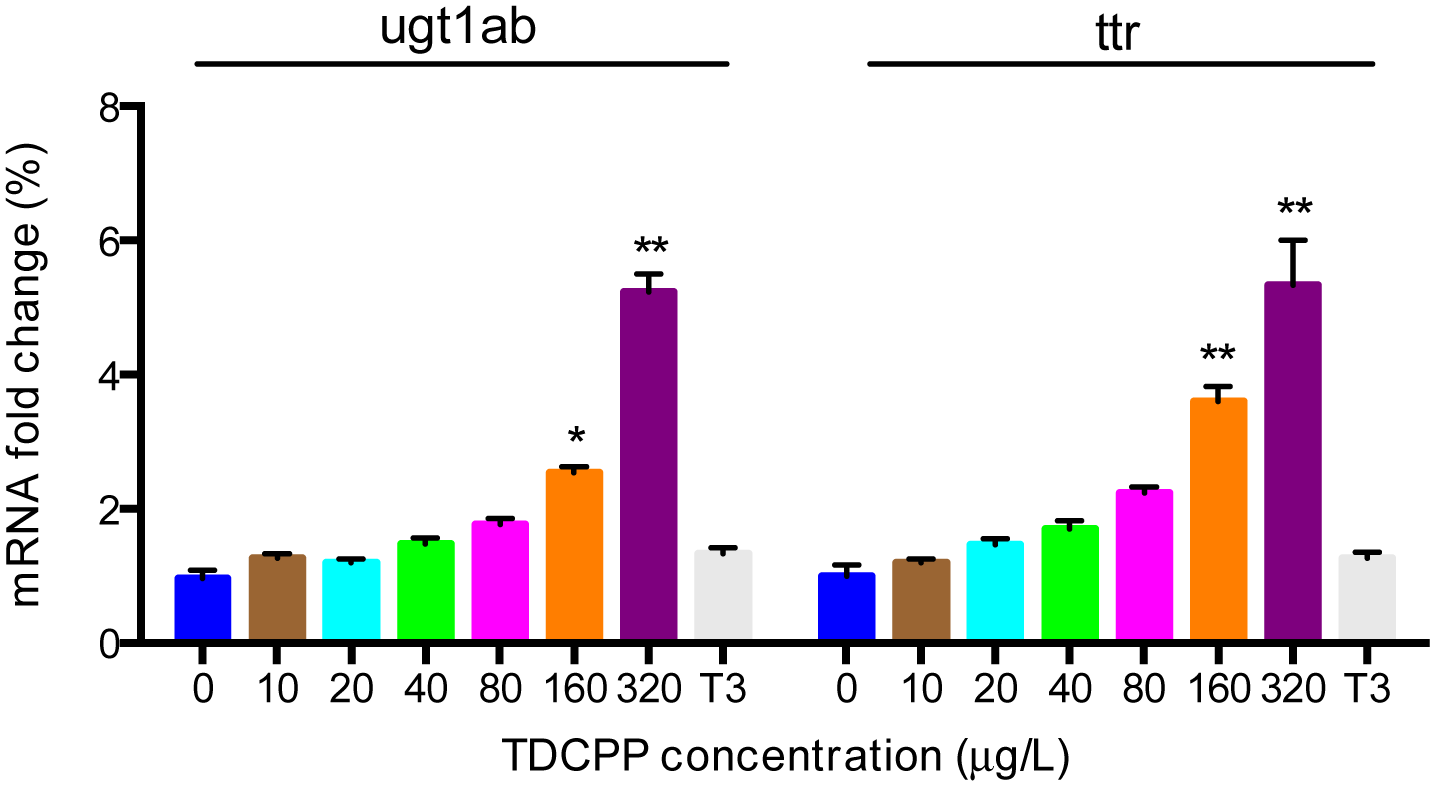
Effects of TDCPP on the expression of ugt1ab and ttr.

### TDCPP treatment increased the T3 and decreased the T4 contents in zebrafish

Exposed to T3 or TDCPP changed the T4 and T3 contents in zebrafish at 5 dpf (Figures 6–7). Tread with T3 or TDCPP increased the T3 contents, while reduced the T4 contents in zebrafish on a dose-dependent manner.

**Figure 6.**
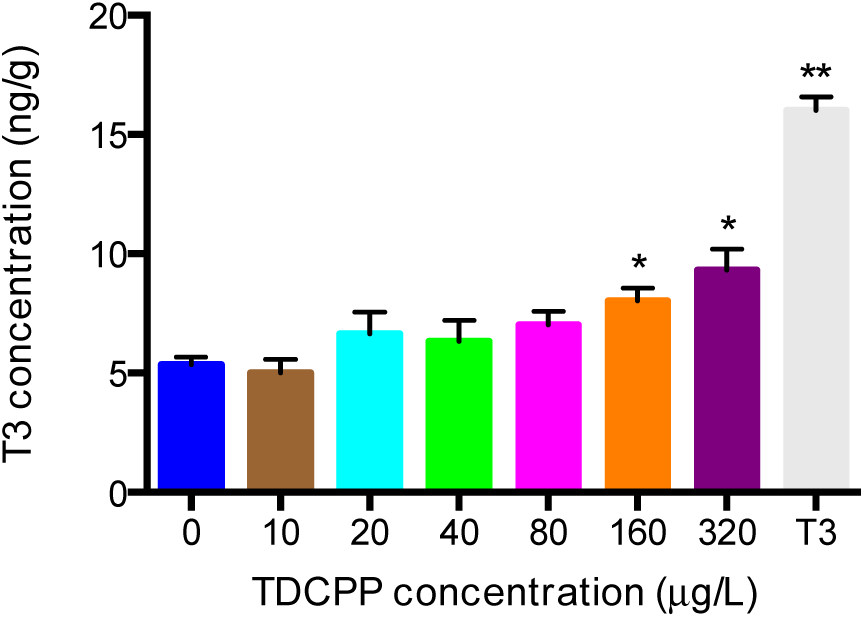
Effects of TDCPP on the T3 concentrations.

**Figure 7.**
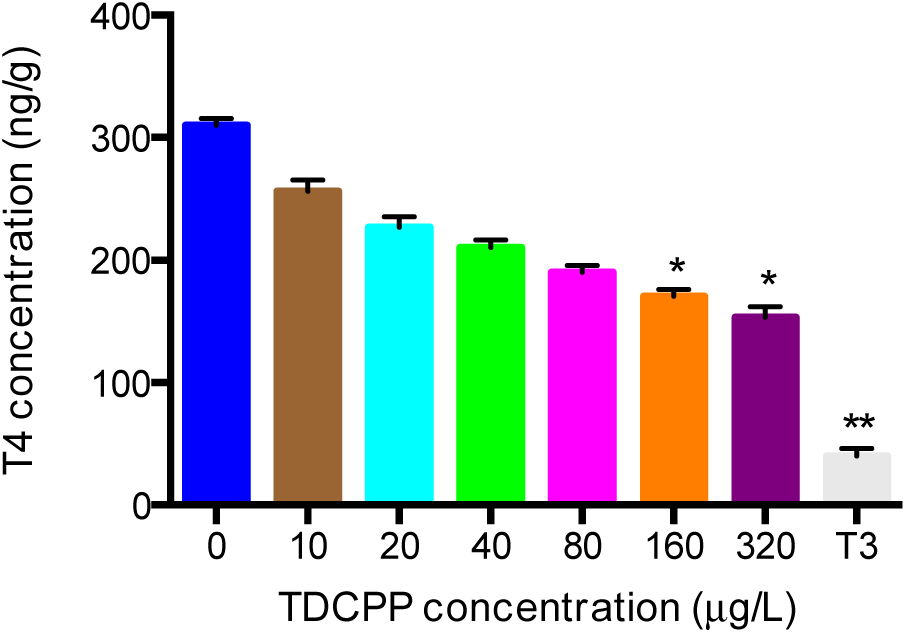
Effects of TDCPP on the T4 contents.

## Discussion

Previous studies report that TDCPP can impair the thyroid endocrine system in zebrafish [6–8, 32]. Our results were in line with these findings that TDCPP could disrupt the thyroid hormone concentrations (T3 and T4) and alter the mRNA expression levels of genes involved in HPT axis of zebrafish. These results confirmed that TDCPP could disrupt the thyroid hormones synthesis, secretion, transport and metabolism [6, 8, 33]. This study showed that zebrafish could be a useful model to monitor the potential effects of drugs on the thyroid endocrine system. Treatment with TDCPP could change the T3 and T4 contents and also the expression of genes involved in HPT axis, which might also affect the reproductive system.

## References

1. Stapleton, H.M., et al., Detection of organophosphate flame retardants in furniture foam and US house dust. Environmental science & technology, 2009. 43(19): p. 7490–7495.

2. Stapleton, H.M., et al., Identification of flame retardants in polyurethane foam collected from baby products. Environmental science & technology, 2011. 45(12): p. 5323–5331.

3. Stapleton, H.M., et al., Novel and high volume use flame retardants in US couches reflective of the 2005 PentaBDE phase out. Environmental science & technology, 2012. 46(24): p. 13432–13439.

4. Dishaw, L.V., et al., Is the PentaBDE replacement, tris (1, 3-dichloro-2-propyl) phosphate (TDCPP), a developmental neurotoxicant? Studies in PC12 cells. Toxicology and applied pharmacology, 2011. 256(3): p. 281–289.

5. Dodson, R.E., et al., After the PBDE phase-out: a broad suite of flame retardants in repeat house dust samples from California. Environmental science & technology, 2012. 46(24): p. 13056–13066.

6. Farhat, A., et al., In ovo effects of two organophosphate flame retardants— TCPP and TDCPP—on pipping success, development, mRNA expression, and thyroid hormone levels in chicken embryos. toxicological sciences, 2013. 134(1): p. 92–102.

7. Liu, X., et al., Effects of TDCPP or TPP on gene transcriptions and hormones of HPG axis, and their consequences on reproduction in adult zebrafish (Danio rerio). Aquatic toxicology, 2013. 134: p. 104–111.

8. Wang, Q., et al., Exposure of zebrafish embryos/larvae to TDCPP alters concentrations of thyroid hormones and transcriptions of genes involved in the hypothalamic–pituitary–thyroid axis. Aquatic toxicology, 2013. 126: p. 207–213.

9. Pedersen, J.A., M. Soliman, and I. Suffet, Human pharmaceuticals, hormones, and personal care product ingredients in runoff from agricultural fields irrigated with treated wastewater. Journal of agricultural and food chemistry, 2005. 53(5): p. 1625–1632.

10. Meeker, J.D. and H.M. Stapleton, House dust concentrations of organophosphate flame retardants in relation to hormone levels and semen quality parameters. Environmental health perspectives, 2010. 118(3): p. 318.

11. Kojima, H., et al., In vitro endocrine disruption potential of organophosphate flame retardants via human nuclear receptors. Toxicology, 2013. 314(1): p. 76–83.

12. Orozco, A. and C. Valverde-R, Thyroid hormone deiodination in fish. Thyroid, 2005. 15(8): p. 799–813.

13. Crump, D., S. Chiu, and S.W. Kennedy, Effects of tris (1, 3-dichloro-2-propyl) phosphate and tris (1-chloropropyl) phosphate on cytotoxicity and mRNA expression in primary cultures of avian hepatocytes and neuronal cells. Toxicological Sciences, 2012: p. kfs015.

14. Costa, L.G., et al., A mechanistic view of polybrominated diphenyl ether (PBDE) developmental neurotoxicity. Toxicology letters, 2014. 230(2): p. 282–294.

15. Blanton, M.L. and J.L. Specker, The hypothalamic-pituitary-thyroid (HPT) axis in fish and its role in fish development and reproduction. Critical reviews in toxicology, 2007. 37(1-2): p. 97–115.

16. Sower, S.A., M. Freamat, and S.I. Kavanaugh, The origins of the vertebrate hypothalamic–pituitary–gonadal (HPG) and hypothalamic–pituitary–thyroid (HPT) endocrine systems: new insights from lampreys. General and comparative endocrinology, 2009. 161(1): p. 20–29.

17. Zoeller, R.T., S.W. Tan, and R.W. Tyl, General background on the hypothalamic-pituitary-thyroid (HPT) axis. Critical reviews in toxicology, 2007. 37(1-2): p. 11–53.

18. Kloas, W., et al., Endocrine disruption in aquatic vertebrates. Annals of the New York Academy of Sciences, 2009. 1163(1): p. 187–200.

19. Zhong, L., et al., Investigation of effect of 17α-ethinylestradiol on vigilin expression using an isolated recombinant antibody. Aquatic toxicology, 2014. 156: p. 1–9.

20. Yang, X., et al., Nucleoporin 62-like protein activates canonical Wnt signaling through facilitating the nuclear import of β-catenin in zebrafish. Molecular and cellular biology, 2015. 35(7): p. 1110–1124.

21. Gu, Q., et al., Generation and characterization of a transgenic zebrafish expressing the reverse tetracycline transactivator. Journal of Genetics and Genomics, 2013. 40(10): p. 523–531.

22. Song, G., et al., Effective gene trapping mediated by Sleeping Beauty transposon. PloS one, 2012. 7(8): p. e44123.

23. Gu, Q., et al., Genetic ablation of solute carrier family 7a3a leads to hepatic steatosis in zebrafish during fasting.Hepatology, 2014. 60(6): p. 1929–1941.

24. Zhai, G., et al., Sept6 is required for ciliogenesis in Kupffer’s vesicle, the pronephros, and the neural tube during early embryonic development. Molecular and cellular biology, 2014. 34(7): p. 1310–1321.

25. Scholz, S., et al., The zebrafish embryo model in environmental risk assessment—applications beyond acute toxicity testing. Environmental Science and Pollution Research, 2008. 15(5): p. 394–404.

26. Parng, C., et al., Zebrafish: a preclinical model for drug screening. Assay and drug development technologies, 2002. 1(1): p. 41–48.

27. Ko, S.-K., et al., Zebrafish as a good vertebrate model for molecular imaging using fluorescent probes. Chemical Society Reviews, 2011. 40(5): p. 2120–2130.

28. Asharani, P., et al., Toxicity of silver nanoparticles in zebrafish models. Nanotechnology, 2008. 19(25): p. 255102.

29. Jopling, C., et al., Zebrafish heart regeneration occurs by cardiomyocyte dedifferentiation and proliferation. Nature, 2010. 464(7288): p. 606–609.

30. Tang, R., et al., Validation of zebrafish (Danio rerio) reference genes for quantitative real-time RT-PCR normalization. Acta biochimica et biophysica Sinica, 2007. 39(5): p. 384–390.

31. Gauthier, K., et al., Different functions for the thyroid hormone receptors TRα and TRβ in the control of thyroid hormone production and post-natal development. The EMBO journal, 1999. 18(3): p. 623–631.

32. Fu, J., et al., Toxicogenomic responses of zebrafish embryos/larvae to tris (1, 3-dichloro-2-propyl) phosphate (TDCPP) reveal possible molecular mechanisms of developmental toxicity. Environmental science & technology, 2013. 47(18): p. 10574–10582.

33. Meng, S., et al., Transdifferentiation Requires iNOS ActivationNovelty and Significance. Circulation Research, 2016. 119(9): p. e129–e138.

